# Engineering a ribozyme with aminoacyl-tRNA synthetase activity

**DOI:** 10.1101/2024.12.12.627959

**Authors:** Gary Gan, Ji Chen, Gerald Manuel, Robert M. Corn, Donald H. Burke, Andrej Luptak, Barbara L. Golden

## Abstract

A ribozyme that can charge a tRNA with amino acids and discriminate between cognate and non-cognate tRNAs is of interest because an RNA with this ability may have been a critical for translation in the transition from the RNA world. In addition, it could provide a tool for incorporating non-canonical amino acids for biotechnology applications. Here, we rationally engineer a ribozyme by fusing a tRNA binding module derived from a T-box riboswitch with a catalytic module (a flexizyme) to generate a ribozyme that can amino acylate a target tRNA. We demonstrate that this ribozyme be readily redesigned to alter tRNA specificity. This ribozyme is compatible with an in vitro translation system and could be used to recode a protein sequence to site-specifically incorporate a non-canonical amino acid.

## Introduction

One of the first events in the evolution of the translation apparatus was likely the emergence of a chemical or enzymatic activity that aminoacylated an RNA. The activity may have been specific to one amino acid and one RNA, or it may have promiscuously generated libraries of charged RNAs that could feed into a primitive translational system. Either way, specificity for both the tRNA and amino acid would need to have emerged for high-fidelity template-directed protein synthesis to arise. The likely existence of such a molecule during the evolution from a primordial RNA world to modern ribosomal protein synthesis suggests that an aminoacylating ribozyme may once have existed and that analogous catalytic RNA molecules could be generated in the laboratory.

Early efforts to generate a ribozyme that could aminoacylate the 3′-terminus of an RNA focused on self-aminoacylating ribozymes. The Yarus lab used in vitro evolution to generate a ribozyme that would self-aminoacylate its 2′- or 3′- terminal hydroxyl groups using an AMP-activated amino acid substrate, similar to that used by the protein tRNA synthetases(Illangasekare, Sanchez et al. 1995). A second ribozyme capable of transferring an AMP-activated amino acid to an internal 2′-hydroxyl was developed by the Famulok group(Jenne and Famulok 1998). Unlike the modern tRNA synthetases, however, these ribozymes are not *trans*-acting catalysts capable of charging tRNAs. Xu et al used the Yarus ribozyme in conjunction with a second ribozyme capable of activating an amino acid substrate by linking it to the phosphate group at its 5′ end. This two-ribozyme system successfully activated an amino acid and transferred it to the 3′-hydroxyl of an RNA substrate (Xu, Appel et al. 2014).

Suga, Szostak and colleagues generated a series of ribozymes that catalyzed the transfer of an amino acid to the termini of RNA molecules. Initially starting with an artificial ribozyme that could transfer an amino acid from the 3′-hydroxyl of a short oligo to its own 5′-hydroxyl terminus(Lee, Bessho et al. 2000), they later developed a programmable ribozyme that could follow this reaction by then transferring the amino acid from its 5ʹ-hydroxyl terminus to the 3ʹ-terminal CCA of a cognate tRNA(Bessho, Hodgson et al. 2002). The identity of the tRNA was determined by base-pairing between the ribozyme and the anti-codon loop of the tRNA. However, this approach provided a maximal yield of only 17% charged tRNA, in large part due to the equilibrium that distributes the amino acid between the charged tRNA product, the 5ʹ end of the ribozyme, and the 3ʹ end of the oligonucleotide substrate.

Suga and colleagues subsequently evolved a series of aminoacylating ribozymes, named flexizymes, by randomizing regions of nucleotides attached to the 5ʹ end of a tRNA (Murakami, Saito et al. 2003, Morimoto, Hayashi et al. 2011). The resulting sequences can be detached from the 5ʹ end of the tRNA, creating a *trans*-acting catalyst. The activity of several of these flexizymes is not restricted by the identity of the amino acid side chain (Murakami, Ohta et al. 2006). Instead, these ribozymes have specificity for the leaving group attached to the carboxyl group of the amino acid substrate. This substrate promiscuity allows the flexizymes to expand the genetic code to include a wide variety of non-biological amino acids *in vitro* (Murakami, Ohta et al. 2006, Ohta, Murakami et al. 2007, Goto, Katoh et al. 2011, Passioura and Suga 2013). Flexizymes, however, bind and act only on the 3′-CCA--terminus of the tRNA substrate. They, therefore, lack the ability to discriminate a cognate tRNA from a set of noncognate tRNAs(Ramaswamy, Saito et al. 2004).

More recently, Ishida et al, evolved a ribozyme, named Tx2.1, capable of charging a specific tRNA substrate with the artificial amino acid N-biotinyl-L-phenylalanine (Ishida, Terasaka et al. 2020). tRNA specificity was obtained using a T-box element. T-box RNAs are naturally occurring riboswitches that can recognize specific tRNAs through interaction with the anticodon loop of the tRNA (Grundy and Henkin 1993, Green, Grundy et al. 2010, Grigg, Chen et al. 2013, Zhang and Ferre-D’Amare 2013, Battaglia, Grigg et al. 2019). By fusing the tRNA-recognizing T-box to a random stretch of nucleotides and carrying out an in vitro selection, a ribozyme capable of aminoacylating the bound tRNA was identified (Ishida, Terasaka et al. 2020).

A ribozyme that has specificity for both the tRNA and amino acid substrates, while interesting from an evolutionary perspective, is also of interest for biotechnology. Such a ribozyme could be designed to charge a tRNA with a non-canonical amino acid (ncAA) or an isotopically-labeled amino acid. If an appropriate anticodon is present on the tRNA, the charged amino acid could subsequently be targeted to a single site or defined set of sites in a growing protein chain to probe enzyme mechanisms, localize proteins within cells, tag proteins for biophysical interrogation, and improve the therapeutic properties of (poly)peptide drugs, among other applications (Wang, Xie et al. 2006, Zhang, Otting et al. 2013, Lang and Chin 2014, Ravikumar, Nadarajan et al. 2015). The Tx2.1 ribozyme was, in fact, used to introduce the artificial amino acid N-biotinyl-L-phenylalanine into a short peptide(Ishida, Terasaka et al. 2020). While this ribozyme provides specificity for the tRNA substrate, its amino acid substrate is limited to the N-biotinyl-L-phenylalanine side chain.

Most ribozymes, like their protein counterparts, are highly specific for their cognate substrates. To obtain alternate specificity, it is necessary to re-engineer or carry out a new *in vitro* selection experiment. The goal of the present study was to generate a ‘universal’ tRNA synthetase that could be rapidly and rationally engineered to recognize alternate tRNA molecules and that could be used with a variety of amino acids. To achieve this, we started with a modular approach that would allow modules to be rapidly switched out to achieve the desired substrate specificity. Specifically, we generated a novel artificial ribozyme, named STARzyme (specific tRNA aminoacylating ribozyme) by fusing a tRNA-recognition module from a T-box RNA with a catalytic module based on the flexizyme (Figure 1A-1C). Creating a ribozyme that can both recognize the anticodon of a tRNA and perform a chemical reaction at the acceptor stem requires the correct orientation of the T-box riboswitch module with respect to the flexizyme module. Through iterative rounds of structure-guided rational design, we demonstrate that a circular permutation of the flexizyme and optimization of the length of the linker and the connector helix generate an aminoacylating ribozyme that exhibits significant tRNA specificity. Furthermore, we demonstrate that tRNA specificity of the STARzyme can be rapidly and rationally reprogrammed by changing the identity of the T-box module.

**Figure 1.**
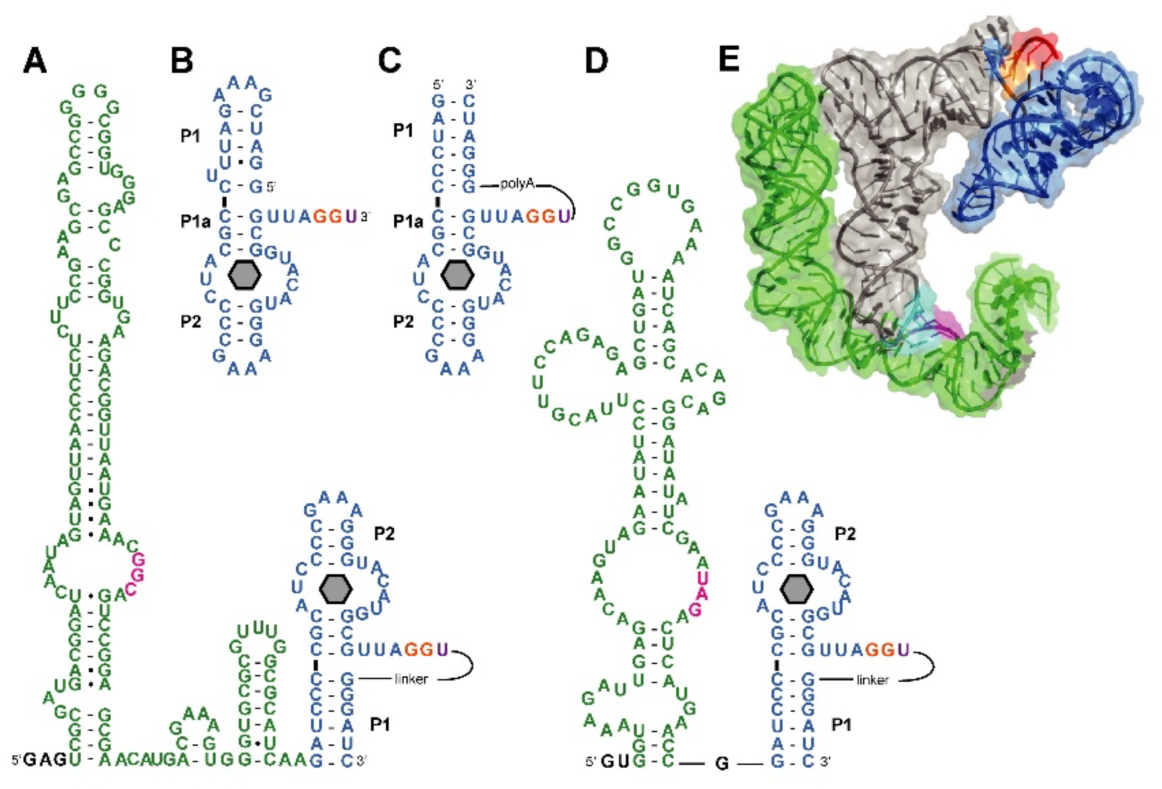
Design strategy for STARzymes. Secondary structures of Gly-STARzyme **(A)** and AMB-STARzyme **(D)**. Green color indicates either *G. kaustophilus glyQS* T-box or *B. subtilus tyrS* T-box; blue color indicates the circularly permuted version of the aminoacylating ribozyme dFx; Magenta nucleotides are involved in anticodon recognition and orange nucleotides bind the CCA tail of the tRNA. The sequence GAG was added to the 5’ end of Gly-STARzyme to facilitate *in vitro* transcription. For AMB-STARzyme, GU was added. **B**. Linear version (original) dFx. The grey hexagon indicates the position of the active site. **C.** Circular permutated version of dFx. **E.** Starting model for tRNA recognition by Gly-STARzyme (PDB ID 4LCK and 3CUL). *glyQS* T-box stem I is shown in green, the bound cognate tRNA (tRNA^Gly^_GCC_) is shown in grey, the Fx3 flexizyme is shown in blue. The CCA tail from the superposed acceptor stem analog in the Fx3 structure is shown in red, while the rest of the acceptor stem analog structure is not shown for simplicity. The specifier and the anticodon are highlighted in cyan and magenta, respectively.

## Methods

### DNA template construction

The template DNA encoding T-box, tRNA, and ribozyme sequences were constructed using overlap PCR. In most cases, the DNA templates were cloned between the *Xba*I and *Hind*III restriction sites of the pUC19 plasmid. The RNA-coding sequences were fused to a T7 RNA polymerase A1 promoter at the 5’ end and 3’ end BsaI restriction site to allow linearization of the plasmid prior to transcription. For the tRNA sequences not starting with a G, a hammerhead self-cleaving ribozyme sequence was inserted between the T7 promoter and the tRNA to facilitate *in vitro* transcription. All DNA templates sequences are listed in Supplementary material Table 1. The WT-DHFR DNA template for protein expression was obtained from the PURExpress *in vitro* protein synthesis kit (NEB). The P105TAG-DHFR DNA template containing a stop codon at position 105 was mutated from the DHFR-WT DNA using the Q5 site-directed mutagenesis kit (NEB).

### RNA sample preparation

RNA samples were prepared by *in vitro* transcription and then purified using denaturing polyacrylamide gel electrophoresis (PAGE), as previously described(Golden 2007). A 50 µl transcription reaction containing 5 µl of 10X transcription buffer, 1 µg of BsaI-linearized DNA template, 2 mM of each NTP and 1 µM of purified T7 RNA polymerase. The transcription reactions were incubated at 37℃ for 1–3 h. For the RNAs cloned downstream of a hammerhead ribozyme, the reaction mixtures were diluted 5 times using 1X transcription buffer after transcription and then incubated at 60℃ for 1 h to enhance ribozyme self-scission. An ethanol precipitation was performed to extract all RNAs from the reaction mixtures and the transcription products were separated through gel electrophoresis. The recovered RNAs were diluted with RNase-free water and concentrated using the 10K centrifugal filters (Merck Millipore) and subsequently stored at −20℃.

### Synthesis of 3,5-dinitrobenzyl ester L-propargyl glycine

All chemicals were purchased from Sigma-Aldrich and used without purification unless otherwise stated. N-boc-L-propargyl glycine was purchased from Chem-Impex. N-boc-L-propargyl glycine (100 mg, 0.469 mmol) and triethylamine (130 μL, 0.938 mmol) were added to 10 mL of dimethylformamide. The mixture was stirred at room temperature until homogenous, then cooled to 0 °C. 3,5-dinitrobenzyl chloride (101 mg, 0.469 mmol) was added to the cooled mixture. The reaction was allowed to warm to room temperature and monitored by thin layer chromatography. After 4 h, the reaction was judged to be complete. 5 mL of diethyl ether were added to the resulting red mixture, and this solution was washed with 0.5 M HCl (5 mL), 4% NaHCO_3_ (5 mL) and with brine (saturated NaCl; 5 mL). The organic phase was dried over Na_2_SO_4_ and concentrated under vacuum. The N-boc protecting group was removed by treatment of the crude product with 2 mL of 4 M HCl in dioxane. After 5 min of stirring at room temperature, a white precipitate formed. The precipitate was isolated and washed with diethyl ether (5 mL) and dried under vacuum to yield the target ester in 81% yield. 1H NMR (600 MHz, d6-DMSO): δ 8.83 (t, J = 1.8, 1H), 8.76 (d, J = 1.2, 2H), 8.65 (br, 3H), 5.51 (q, J = 13.8, 2H), 4.44 (t, J = 5.4, 1H), 3.32 (t, J = 2.4, 1H), 2.91 (m, 1H), 2.837 (m, 1H). ESI-MS: m/z calculated for C_12_H_11_N_3_O_6_ (M+Cl)− 328.065, found 328.100.

### Gel shift assay

To refold RNAs, 40 µM ribozyme and 10 µM tRNA were heated separately at 95℃ for 2 min and then slowly cooled to room temperature over 10 min in the refolding buffer containing 50 mM HEPES-KOH (pH 7.5) and 100 mM KCl. 20 mM MgCl_2_ was added into ribozyme and tRNA followed by 10 min incubation at room temperature. A total volume of 20 µl was set for the binding reaction containing 4 µl of 5X reaction buffer (250 mM HEPES-KOH pH 7.5, 500 mM KCl and 50 mM MgCl_2_), 2 µl of 10 µM tRNAs, 4 µl of 40 µM ribozymes to make a final concentration up to 8 µM and sufficient water to make a final volume 20 µl. The binding reactions were incubated at room temperature for 1 h followed by the addition of 5 µl of 50% glycerol. Five µl of each sample were loaded onto a 6% native PAGE gel and then fractionated at 4℃ for 1.5 h with a 10 W power setting. The gel was stained with Stains-all (Sigma) and scanned with a Bio-Rad image system.

### *In vitro* aminoacylation assay

For each reaction, 10 µl of 10 µM ribozyme and 10 µl of 2 µM tRNA were first refolded separately as described above. The ribozyme was then mixed with tRNA and the mixture was incubated at room temperature for 1 h followed by 4℃ for 10 min. The aminoacylation was initiated by the addition of 1 µl of 100 mM DBE-propargyl glycine dissolved in DMSO and the reaction was incubated at 4℃ for up to 6 h. At each time-point, 2 µl of the mixture was removed and mixed with 8 µl of the loading buffer containing 100 mM sodium acetate (pH 5.2), 7 M urea, 0.05% bromophenol blue and 10 mM EDTA. The above mixture was loaded to a pre-run 8% acidic denaturing PAGE gel containing 7 M urea and 100 mM sodium acetate (pH 5.2) and fractionated at 10 W for up to 19 h at 4℃. The low pH of the running buffer prevents the hydrolysis of aminoacylated tRNA. The gel was stained with SYBR GREEN II (Invitrogen) and scanned by Typhoon FLA 9500 (GE Healthcare). Under these conditions, the band intensity of the SYBR GREEN II stained gel has a linear relationship with the amount of total *E. coli* tRNA ranging from 0 to 0.14 µg **(Figure S7)**.

The aminoacylation of total *E. coli* tRNA was performed using the same method, but the reactions contained 10 µl of 20 µM ribozyme and 10 µl of 0.1 µg/µl (about 4 µM) total *E. coli* tRNA (Sigma). Two µl of the reaction product was mixed with the loading buffer and loaded onto a gel containing what gel electrophoresis and analysis for how long. Three major bands corresponding to a mixture of 86 tRNAs can be observed on gel and the bottom band (band 3) was chosen to represent the total *E. coli* tRNA because the decreasing of its band intensity can be reasonably explained as the increasing of aminoacylation.

### Kinetics assay

For single-turnover kinetics, 10 µM AMB-STARzyme and 2 µM tRNA were used in the aminoacylation reaction. 2 µl of the reaction product was used for PAGE and the intensity of charged and uncharged bands was quantified. The data were fit to a single-exponential equation (Synergy KaleidaGraph)

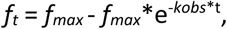

where *f_t_* is the fraction of aminoacylated tRNA at time t, *f_max_* is the fraction of aminoacylated tRNA extrapolated to infinite time and *k_obs_* is the observed rate constant.

For multiple-turnover kinetics, reactions were performed with a fixed concentration of tRNA (1 μM) and various concentrations of AMB-STARzyme (0.1, 0.2, 0.5, 1 μM). The charged tRNAs were separated from the uncharged tRNAs, and the fraction of aminoacylated tRNA were calculated and converted into the concentration of aminoacylated tRNA (μM). The turnover numbers were equal to the results of aminoacylated tRNA (μM)/AMB-STARzyme (μM).

### *In vitro* translation

PURExpress ΔRF123 kits (NEB) were used for *in vitro* translation reactions, with a total reaction volume of 30 µl for each sample. The components were added to a tube in the following order: 10 µl of solution A (containing tRNAs, amino acids, and rNTPs), 7.5 µl of solution B (containing ribosome, T7 RNA polymerase, translation factors, and tRNA synthetases), 0.5 µl of release factor 2, 0.5 µl of release factor 3, 0.5 µl of RNase inhibitor (NEB), 0.5 µl of 360 mM magnesium acetate, 0.5 µl of 120 mM ATP, 1 µl of 100 mM DBE- propargyl glycine dissolved in DMSO, 2 µl of 400 µM *M. jannaschii* tRNA^AMB^_CUA_, 6 µl of 330 µM AMB-STARzyme, 0.5 µl of RNase-free water and 0.5 µl of 600 ng/µl P105TAG-DHFR DNA template to make a final volume 30 µl. The reactions were incubated at 37℃ for 3 h.

### Peptide preparation

Protein products from *in vitro* translation (30 µl) were separated by SDS-PAGE gel electrophoresis and stained with Coomassie blue. The target bands were excised and in-gel digested for proteomics analysis. Target bands were treated with 100 µl of dithiothreitol 10 mM for reduction and 100 µl of 40 mM iodoacetamide for alkylation. Samples were dried in vacuum centrifuge followed by the addition of 1 µl of 0.2 µg/µl trypsin (Sigma) in 50 mM TEAB (triethylammonium bicarbonate) and 99 µl of 50 mM TEAB for overnight digestion at 37℃. The tryptic peptides were then desalted and dissolved in 20 µl of buffer containing 0.1% formic acid in 80% acetonitrile for further analysis.

### Mass spectrometry analysis

For MALDI-TOF analysis, 1 µl of the peptide samples from peptide preparation was mixed with 1 µl of 10 mg/ml CHCA (α-cyano-4-hydroxycinnamic acid) solution and spotted on MALDI plate. Positive ion mass spectra were obtained on an Applied Biosystems/MDS SCIEX 4800 MALDI TOF/TOF (Thermo Fisher Scientific) in the reflector mode, and the mass range for scanning was 500-4000 amu.

For LC-MS/MS analysis, the peptide samples from peptide preparation were dried in vacuum centrifuge and dissolved in 10 µl of buffer containing 0.3% formic acid in 3% acetonitrile. 4 µl of the resuspended samples were injected into EASY-nLC 1000 system coupled with a Velos Pro LTQ-Orbitrap mass spectrometer (Thermo Fisher Scientific) for analysis. Peptides were separated on a 45 cm C18 resin column. The mobile phase buffer consisted of 0.1% formic acid in ultrapure water and the elution buffer consisted of 0.1% formic acid in 80% acetonitrile. A linear gradient of elution buffer from 6% to 30% was applied over 60 min at a flow rate of 250 nL/min to separate the peptides. The eluent was introduced into the mass spectrometer with the data-dependent mode setting, and a full-scan MS was conducted from m/z 350 to 1500 (the resolution of 30000 at m/z = 400). The target precursor ions were selected for a MS/MS analysis using a normalized collision energy.

## Results

The goal of this study was to rationally engineer a ribozyme that could amino acylate a target tRNA and be readily redesigned to alter tRNA or amino acid specificity. Our strategy was to link a readily available tRNA-binding module to a previously described catalytic module. For the tRNA-binding component, we started with the tRNA^Gly^_GCC_ binding T-box (*glyQS* T-box) RNA from *Geobacillus kaustophilus*(Gagnon, Breton et al. 1994, Niwa, Yamagishi et al. 2009) (shown in green in Figure 1). The *glyQS* T-box was chosen because it has a relatively minimal structure, it is thermodynamically stable, and its structure is well characterized (Grigg, Chen et al. 2013, Grigg and Ke 2013, Zhang and Ferre-D’Amare 2013, Li, Su et al. 2019). For the catalytic module, we used a flexizyme variant (dFx) specific for amino acids activated by a dinitrobenzyl ester (DBE). dFX is an artificial ribozyme that accepts a wide variety of nonaromatic amino acid substrates (Figure 1B)(Murakami, Ohta et al. 2006). We generated a three-dimensional model of our fusion ribozyme using the crystal structure of the T-box riboswitch bound to a cognate tRNA (Grigg and Ke 2013, Zhang and Ferre-D’Amare 2013). We then superposed the acceptor stem of the bound tRNA with the acceptor stem analog present in the crystal structure of the Fx3 flexizyme (Xiao, Edwards et al. 2008) (shown in blue in Figure 1). Fx3 is a flexizyme variant that is similar in structure to dFx and provides a model for the size and orientation of dFx bound to the CCA tail of the tRNA. This model, shown in Figure 1D, demonstrates that the T-box riboswitch and the flexizyme should not sterically interfere with each other when docked to a tRNA and could potentially be used to engineer a ribozyme with tRNA synthetase activity.

### Circular permutation of dFx is required for catalytic activity

We first explored the catalytic activity of a linear fusion of the T-box RNA and the dFx flexizyme. To measure the catalytic activity of the ribozymes, we used an activated amino acid substrate containing a propargyl glycine group (Figure S1A) activated by a 3,5-dinitrobenzyl ester. Reactions were performed at 4 °C to minimize hydrolysis of the substrate (Supporting Information, Figure S2). At this temperature, the half-life of the substrate is over 40 h.

The linear fusion ribozyme was catalytically active; however, its activity was lower than that observed for isolated dFx and it did not distinguish between the cognate tRNA and the noncognate one (data not shown). The tRNA-binding properties (Supporting Information, Figure S3) and the presence of catalytic activity of the linear fusion suggested that the T-box and flexizyme modules are both folded and functional. However, the lack of discrimination and diminished catalytic activity suggest that the tRNA substrate cannot simultaneously bind the flexizyme and T-box modules in this initial design.

The working three-dimensional model of the fusion ribozyme (Figure 1D) suggested that a circular permutation of the dFx flexizyme created by linking the 5′ and 3′ ends and opening the P1 loop (Figure 1C) might be a better topology for fusing the two modules. It was unclear, however, how best to link the 5′ and 3′ termini of dFx because the CCA-binding site is located on a single-stranded tail at the 3′ end of dFx and disrupting the structure in this region would therefore be detrimental to ribozyme activity. To provide some flexibility, a number of adenosines were used to link the 5′ end of the parental dFx molecule with the 3′ end containing the CCA-binding site (Figure 1C). This insertion was not predicted to interfere with formation of the native secondary structure of dFx ribozyme. The 3′ end of the *glyQS* T-box was connected to the new 5′ end of the circularly permuted dFx RNA (Figure 1A). This fusion ribozyme was named the specific tRNA aminoacylating ribozyme, or ‘STARzyme.’ The overall topology of STARzyme resembles that of the wild-type *glyQS* T-box (Yousef, Grundy et al. 2003), where the aptamer domain is connected to a helix P1 that is then linked to an antiterminator containing the binding motif for the CCA tail.

STARzyme variants with seven, eight, or ten adenosine linkers in the circularly permuted dFx module were analyzed for catalytic activities in the presence of either cognate or noncognate tRNAs. The aminoacylation activity was monitored by a gel mobility assay (Figure 2A). Maximal activity of the flexizyme requires base pairing between the dFx module and the discriminator nucleotide that precedes the CCA tail of the tRNA (Murakami, Saito et al. 2003) and Figure S1. Thus, tRNA^Gly^_GCC_ U73A mutant is considered as cognate because it can base pair with both the specifier nucleotides of T-box and U46 of dFx. The STARzyme with an eight-nucleotide linker (STAR-A8) was almost as active as isolated dFx in the presence of its cognate tRNA^Gly^_GCC_ U73A. The observed rate constant for STARzyme was 0.21 ± 0.02 h^-1^, whereas the isolated dFx exhibits a rate constant of 0.54 ± 0.08 h^-1^. In contrast, STAR-A8 was ∼4-fold less active in the presence of the noncognate tRNA^Ile^_GAU_ (k_obs_ = 0.055 ± 0.0032 h^-1^) than in the presence of the cognate tRNA^Gly^_GCC_ U73A (Figure 2A, Table 1). The eight-nucleotide linker proved optimal, because the STARzyme with a ten-nucleotide linker displayed less discrimination, with only a ∼2-fold change in the observed rate constants for cognate and non-cognate tRNA (k_obs_ = 0.17 ± 0.05 h^-1^ for the cognate tRNA^Gly^_GCC_ U73A *versus* k_obs_ = 0.090 ± 0.029 h^-1^ for tRNA^Ile^_GAU_), whereas the STARzyme with a seven nucleotide linker was lower in catalytic activity (k_obs_ = 0.092 ± 0.015 h^-1^ for the cognate tRNA versus k_obs_ = 0.026 ± 0.006 h^-1^ for tRNA^Ile^_GAU_; Table 1).

**Figure 2.**
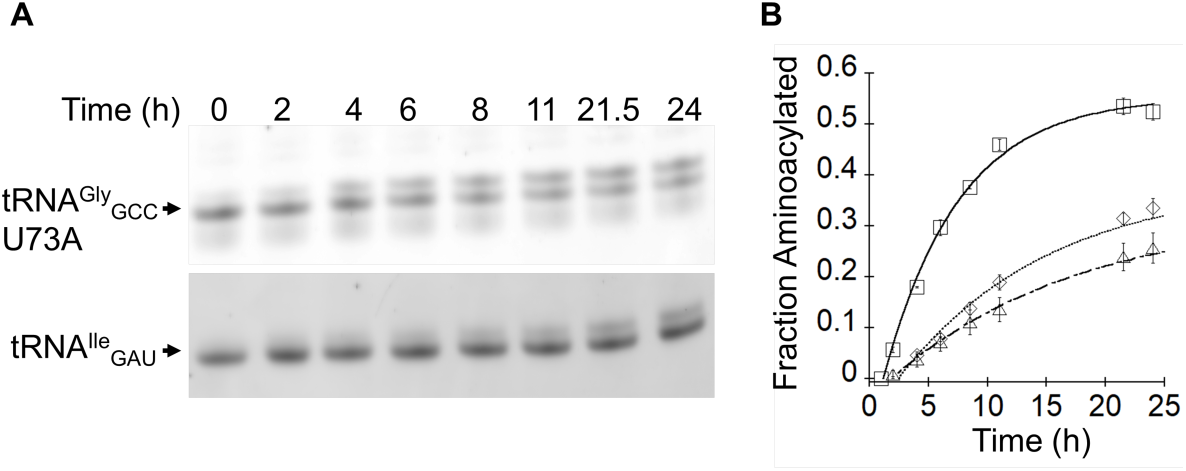
Testing the catalytic activity of Gly-STARzyme against cognate and noncognate tRNAs. **A.** Representative gel showing the Gly-STARzyme-mediated aminoacylation reaction over time in the presence of the cognate *G. kau* tRNA^Gly^_GCC_ U73A (top) or the noncognate *G. kau* tRNA^Ile^_GAU_ (bottom). **B.** Reaction progress curves for Gly-STARzyme against *G. kau* tRNA^Gly^_GCC_ U73A (open squares), *G. kau* tRNA^Gly^_GCC_ (open diamonds) or *G. kau* tRNA^Ile^_GAU_ (open triangles).

**Table 1.**
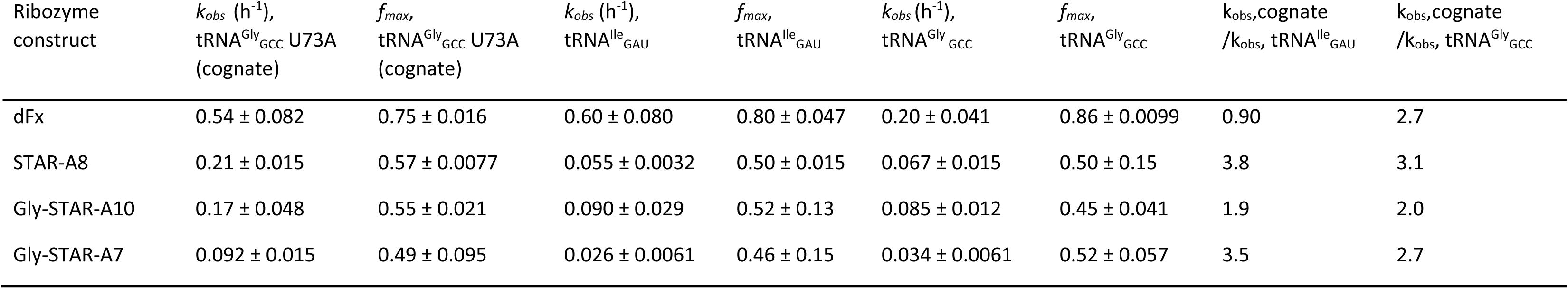
Kinetic parameters for dFx and Gly-STARzymes. Parameters reported here represent average values from three independent measurements. Errors stand for standard deviations.

The difference in STARzyme activity between cognate tRNA^Gly^_GCC_ U73A and non-cognate tRNA^Ile^_GAU_ suggests that the fusion ribozyme is using the specifier nucleotides in the T-box module to recognize the anticodon loop and thereby gain specificity. To test whether the fusion was using the discriminator base on tRNA as an additional specificity determinant, the reaction with wild-type tRNA^Gly^_GCC_ was also characterized. As observed for the reaction of the dFx ribozyme(Murakami, Saito et al. 2003), the STARzyme was 2- to 3- fold less active with the wild-type tRNA^Gly^_GCC_ than the tRNA^Gly^_GCC_ U73A mutant (Figure 2B, Table 1). Therefore, STARzyme likely achieves tRNA discrimination by recognizing both the anticodon and the discriminator base of the acceptor stem.

### Detection of the tRNA-linked amino acid

Our activity assay uses a gel mobility assay previously shown to be appropriate for monitoring the aminoacylation of tRNA (Hamashima, Fujishima et al. 2011). To establish unambiguously that the shift in gel mobility is due to addition of the ncAA to the tRNA, we took advantage of the alkynyl group in the ncAA, which allows covalent modification through copper-catalyzed ‘click’ cycloaddition(Kolb, Finn et al. 2001). tRNA^Gly^ U73A was aminoacylated using either the dFx flexizyme or the STARzyme for a period expected to provide the maximal yield of tRNA product. The reaction mixture was treated with Alexa Fluor 488 picolyl azide, which consists of a fluorophore and an azide group, and thus should specifically label the alkyne group on the aminoacylated RNAs but not the ribozyme or unreacted tRNA. Reactions and controls were loaded side-by-side into duplicate lanes on a single polyacrylamide gel. After electrophoresis, the gel was cut into two halves. The first half was stained with SYBR green II to visualize all RNAs, and the second half remained unstained to allow detection of the fluorescent signal from the Alexa Fluor 488 probe. An Alexa Fluor-labeled band was observed at the molecular weight corresponding to the aminoacylated tRNA product only when all of the required components of the aminoacylation reaction were present in the mixture (Figure 3). When small molecules were removed from the reaction by ultrafiltration subsequent to the click reaction, we observed only a single fluorescently-labeled band, again corresponding to the size of the supershifted aminoacylated tRNA product. Thus, the STARzyme is catalyzing the addition of the artificial amino acid to the tRNA substrate.

**Figure 3.**
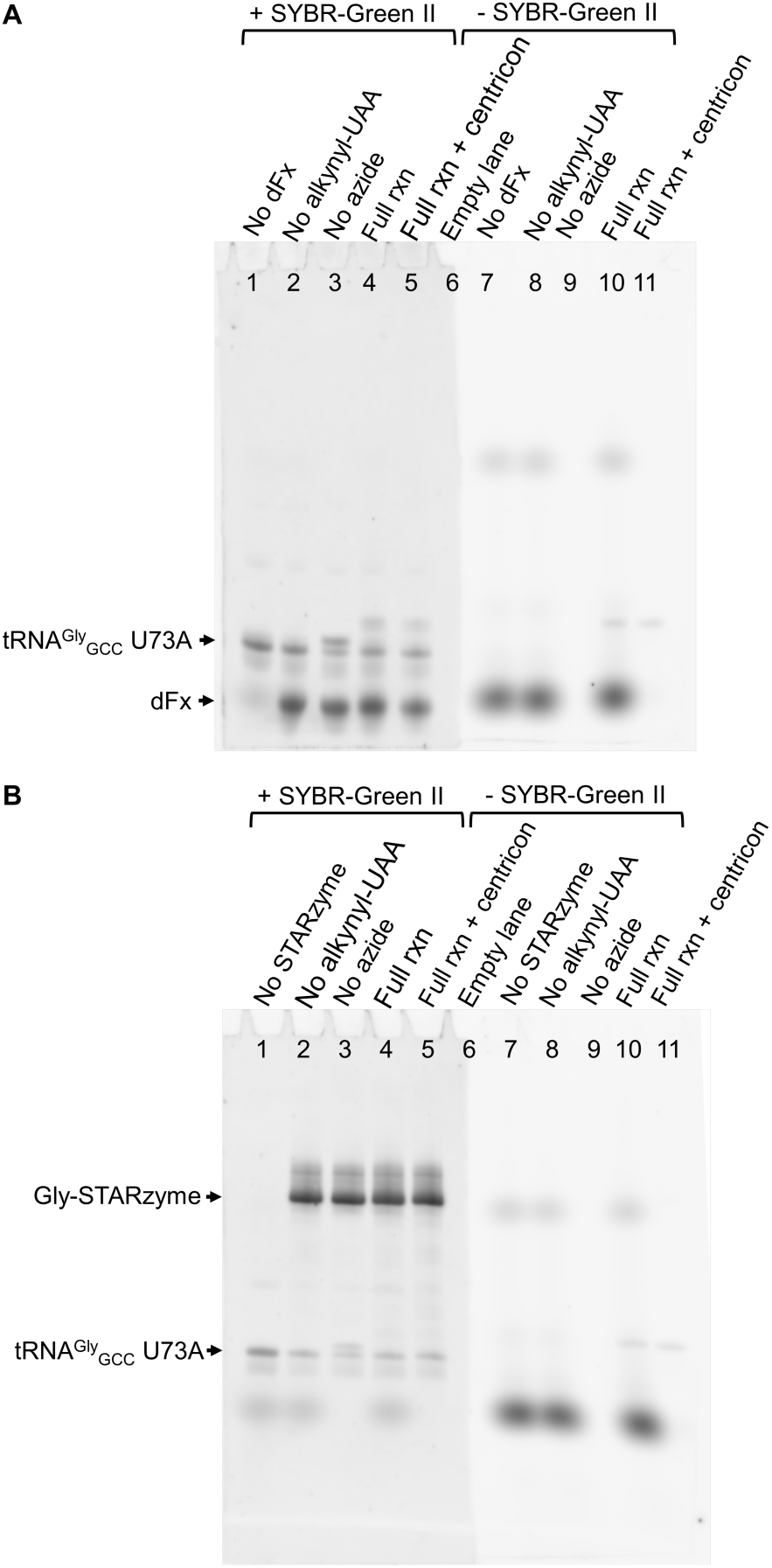
Identification of the aminoacylation reaction product using click chemistry. **A.** Aminoacylation assays using dFx. The aminoacylation reactions were split and separated on the same gel. The left half of the gel was stained with SYBR-Green II, while the right portion of the gel was unstained. The two half gels were placed side by side prior to scanning. The aminoacylation reactions and click reactions were performed in all lanes as described in Method and Materials, except for intentional omission of the indicated reaction components. Full rxn + centricon: same as “Full rxn”, except that the reaction sample was passed through an Amicon Ultra centrifugal filter unit with a 10K NMWL cutoff to remove unreacted fluorophore. In the SYBR-Green II stained gel, the putative charged tRNA is supershifted upon the addition of Alexa Fluor 488-tagged picolyl azide. The identity of the supershifted band was further confirmed by detection of Alexa Fluor 488 fluorescence. Upon ultrafiltration, the single fluorescent band (lane #11) on the right half of the gel comigrates with the “supershifted” band on the left half of the gel (lane #4 or lane #5). **B.** Same as panel A except that Gly-STARzyme was used instead of dFx.

### Rational engineering of tRNA specificity

To demonstrate that we could rationally reprogram the tRNA specificity, we replaced the *glyQS* T-box of the first generation STARzyme with the *Bacillus subtilis tyrS* T-box. We chose the *B. subtilis tyrS* T-box because it had been previously demonstrated that its specificity for the tRNA anticodon sequence could be rationally engineered. In particular, mutating the specifier nucleotides of the T-box that base-pair with the anticodon from UAC to UAG was demonstrated to shift the tRNA binding specificity from wild-type tRNA^Tyr^_GUA_ to the amber suppressor tRNA^AMB^_CUA_ (Grundy, Hodil et al. 1997, Grundy, Collins et al. 2000). To facilitate the correct folding *in vitro*, a truncated *tyrS* T-box spanning stem I and lacking the pseudoknot element was used for the second generation STARzyme. This truncated T-box folds efficiently *in vitro* and is predicted to be sufficient for specific and high-affinity tRNA binding. The mutant *tyrS* T-box stem I (AMB-T-box) was fused with circularly permuted dFx as described for the first generation STARzyme.

Activity assays were conducted with four tRNAs: the wild-type tRNA^Tyr^_GUA_ and amber suppressor tRNA^AMB^_CUA_ from *B. subtilis* and the corresponding tRNA^Tyr^_GUA_ and tRNA^AMB^_CUA_ from *E. coli.* The *E. coli* tRNAs were included both to define the tRNA structural elements that contribute to reprogrammability and as preliminary evaluation of the suitability of AMB-STARzyme for use in bacteria. Aminoacylation reactions were performed using the conditions established above, and the products were separated by denaturing PAGE. As shown in **Figure 4**, the slower-migrating bands can be clearly observed over time only in the group of *B. sub* tRNA^AMB^_CUA_ and *E. coli* tRNA^AMB^_CUA_, indicating that both amber suppressor tRNAs were efficiently aminoacylated. No slower-migrating bands can be observed in the control group of *B. sub* tRNA^Tyr^_GUA_ and *E. coli* tRNA^Tyr^_GUA_. Considering that there is only one nucleotide difference (**<u>C</u>**UA and **<u>G</u>**UA, at the anticodon loop) between the cognate and noncognate tRNAs, the above results suggested that AMB-STARzyme has a high degree of specificity. Another two noncognate tRNAs (*G. kau* tRNA^Gly^_GCC_ and *G. kau* tRNA^Ile^_GAU_) were also tested and they cannot be efficiently aminoacylated by AMB-STARzyme **(Figure S2)**. These results demonstrated that the AMB-T-box and dFx modules are functional in AMB-STARzyme and yield a highly specific ribozyme capable of aminoacylating suppressor tRNAs with potential applications in Amber-suppression recoding.

**Figure 4.**
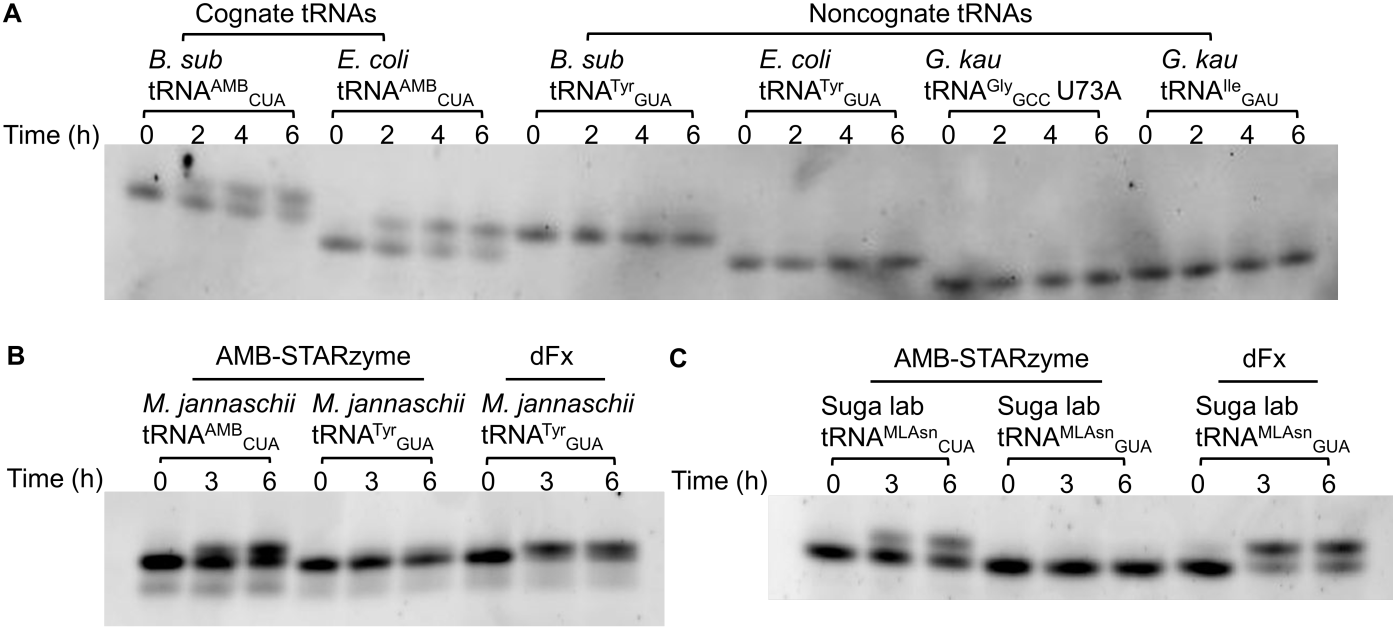
Catalytic activity of AMB-STARzyme against cognate, noncognate and orthogonal tRNAs. **A.** The gel showing AMB-STARzyme-mediated aminoacylation reaction over time in the presence of cognate tRNAs (*B. sub* tRNA^AMB^_CUA_, *E. coli* tRNA^AMB^_CUA_) and noncognate tRNAs (*B. sub* tRNA^Tyr^_GUA_, *E. coli* tRNA^Tyr^_GUA_, *G. kau* tRNA^Gly^_GCC_ U73A, *G. kau* tRNA^Ile^_GAU_). **B.** The gel showing the orthogonal *M. jannaschii* tRNA^AMB^_CUA_ aminoacylated by AMB-STARzyme, and its mutant *M. jannaschii* tRNA^Tyr^_GUA_ aminoacylated by AMB-STARzyme and dFx. **C.** Suga lab tRNA^MLAsn^_CUA_ and its mutant tRNA^MLAsn^_GUA_, were tested under the same conditions as in panel B.

Although both amber suppressor tRNAs *B. sub* tRNA^AMB^_CUA_ and *E. coli* tRNA^AMB^_CUA_ can be efficiently aminoacylated by the AMB-STARzyme, neither is appropriate for use in an *E. coli in vitro* translation system because they are not orthogonal and can be mischarged by endogenous *E. coli* aaRS(Himeno, Hasegawa et al. 1990). In contrast, archaeal tRNAs have been demonstrated to be orthogonal in *E. coli* system as they are poor substrates for *E. coli* tRNA aaRS but function efficiently in protein translation(Wang, Brock et al. 2001, Gundllapalli, Ambrogelly et al. 2008). In the *E. coli* system, a commonly used archaeal amber suppressor tRNA is *M. jannaschii* tRNA^AMB^_CUA_ derived from a tyrosine tRNA. Artificial and engineered tRNAs have also been developed as orthogonal tRNAs for use in *E. coli* systems. In flexizyme research from the Suga group, three artificial tRNAs with unique body sequences were successfully applied in *E. coli* system and a high orthogonality of these tRNAs was observed(Gundllapalli, Ambrogelly et al. 2008). For STARzyme characterization, we focused on the *M. jannaschii* and the Suga group tRNAs and altered their anticodon to generate CUA amber suppressor tRNAs.

STARzyme carries three elements that can serve as tRNA recognition determinants. One results from base-pairing of the anticodon with the specificity nucleotides of the T-box. An additional specificity determinant is the discriminator nucleotide at position 73 on tRNA. Previous studies demonstrated that aminoacylation was most productive when the discriminator base at position 73 is A, which is complementary to the 3’-terminal U in the Fx3 or dFx ribozyme(Murakami, Saito et al. 2003). Thus, to achieve the maximal activity of the AMB-STARzyme, all tRNAs used in this research have a naturally occurring A73 or an A73 variant was introduced. The secondary structure of AMB-STARzyme and tRNAs are shown in **Figure 1** and **Figure S1**. Finally, there is also a quaternary interaction between the elbow region of the tRNA and the T-box module. The archaeal tRNAs and artificial RNAs should preserve the first two of these, but they have significantly different sequence in the elbow region that could potentially interfere with recognition by the *B. subtilis* AMB-T-box module. Nevertheless, four out of six of the amber suppressor tRNAs were efficiently aminoacylated by this STARzyme (**Figure 4**). To rule out the possibility that the STARzyme’s circularly permuted flexizyme module might aminoacylate these tRNAs independent of binding to the T-box module, we mutated the anticodon loops of three representative tRNAs (highlighted) to GUA (corresponding to anticodon of tyrosine tRNAs) and tested them again. As expected, none of the mutants was efficiently aminoacylated by AMB-STARzyme. In contrast, all mutants were efficiently aminoacylated by dFx **(Figure S3)**. We chose the *M. jannaschii* tRNA^AMB^_CUA_ for further characterization, as this tRNA has been widely used in numerous reports because of its high orthogonality and suppression efficiency(Wan, Tharp et al. 2014).

### Kinetic Parameters of AMB-STARzyme

Single- and multiple-turnover kinetic parameters of AMB-STARzyme were evaluated separately. To measure the single-turnover rate, kinetic experiments were performed over the course of 12 h as described above using 1 μM cognate or noncognate tRNAs were mixed with 5 μM AMB-STARzyme in the presence of 5 mM DBE-propargyl glycine. Apparent rate constant was obtained by fitting the fraction of aminoacylated tRNA versus time using the first-order kinetic equation (**Fig. 5**). Given that the flexizyme modules evolved under single-turnover conditions and Fx3, a prototype flexizyme, was reported to display a maximum turnover number of 14 (Murakami, Saito et al. 2003). We anticipated that the STARzyme may be limited in its ability to aminoacylate multiple tRNAs during the course of the reactions. We therefore carried out the experiments under multiple turnover conditions using a constant tRNA (1 μM) concentration and four different concentrations of AMB-STARzyme (0.1, 0.2, 0.5, 1 μM). The results indicate that less than one tRNA/STARzyme molecule are aminoacylated over the course of 24 h (Figure S4), suggest that AMB-STARzyme does not exhibit significant multiple turnover under these conditions.

**Figure 5.**
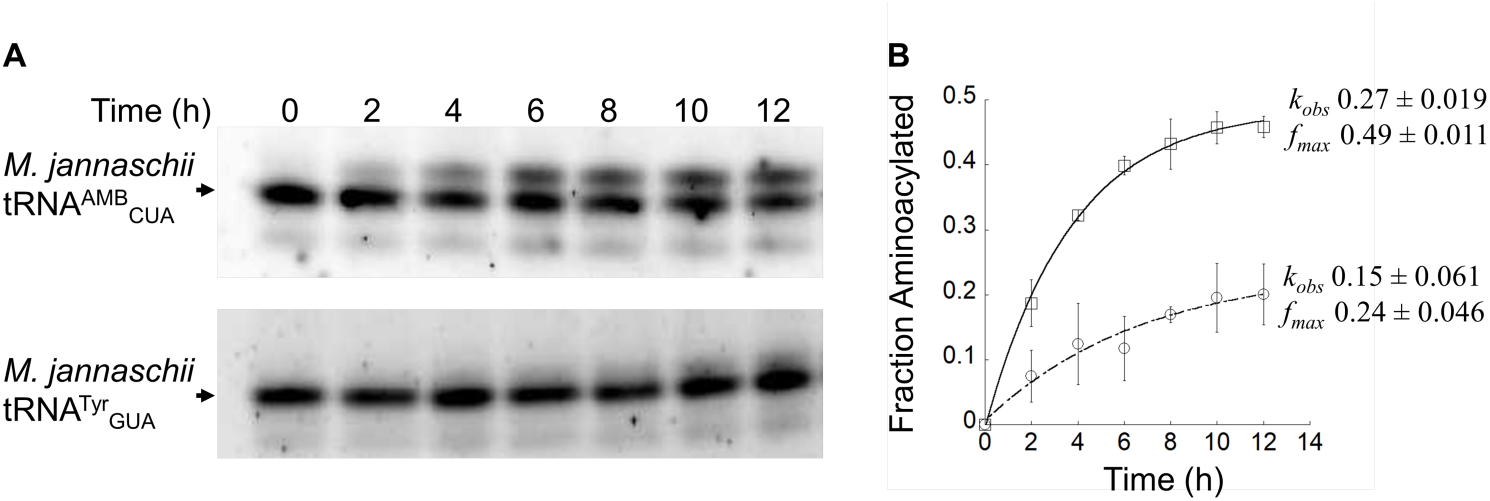
Kinetic assay of AMB-STARzyme against *M. jannaschii* tRNA^AMB^_CUA_ and tRNA^Tyr^_GUA_. **A.** The gel showing AMB-STARzyme-mediated aminoacylation reaction over time in the presence of *M. jannaschii* tRNA^AMB^_CUA_ (top) or tRNA^Tyr^_GUA_ (bottom). **B.** Reaction progress curves for AMB-STARzyme against *M. jannaschii* tRNA^AMB^_CUA_ (open square) and tRNA^Tyr^_GUA_ (open circle).

### Orthogonality of AMB-STARzyme

Aminoacylation of *M. jannaschii* tRNA^AMB^_CUA_ by AMB-STARzyme suggests this ribozyme has the potential to be useful in generating substrates for translation. However, for use in an *E. coli in vitro* translation system, both the AMB-STARzyme and the tRNA substrate must be orthogonal to the components in the translation system. The orthogonality of an aaRS is defined by two parts: orthogonality to amino acid and orthogonality to tRNA. Because the STARzyme requires an activated amino acid with an artificial DBE leaving group, it is not expected to react with any of the endogenous amino acids. Prior studies have established that *M. jannaschii* tRNA^AMB^_CUA_ is not charged to a significant degree by endogenous *E. coli* aaRSs(Hong, Ntai et al. 2014). Therefore, the main concern was whether the AMB-STARzyme mischarges any of the 86 endogenous *E. coli* tRNAs. To test the orthogonality of AMB-STARzyme with respect to endogenous *E. coli* tRNAs, the *in vitro* aminoacylation reactions were performed using the total *E. coli* tRNA (Sigma) and AMB-STARzyme. dFx was used as a positive control. The results indicated that a maximum of ∼ 3% of the total *E. coli* tRNA was charged by AMB-STARzyme in a 12-h incubation, while the number for dFx is 28% (**Figure 6**). Thus, the T-box module of the AMB-STARzyme appears to bestow significant specificity for the target tRNA.

**Figure 6.**
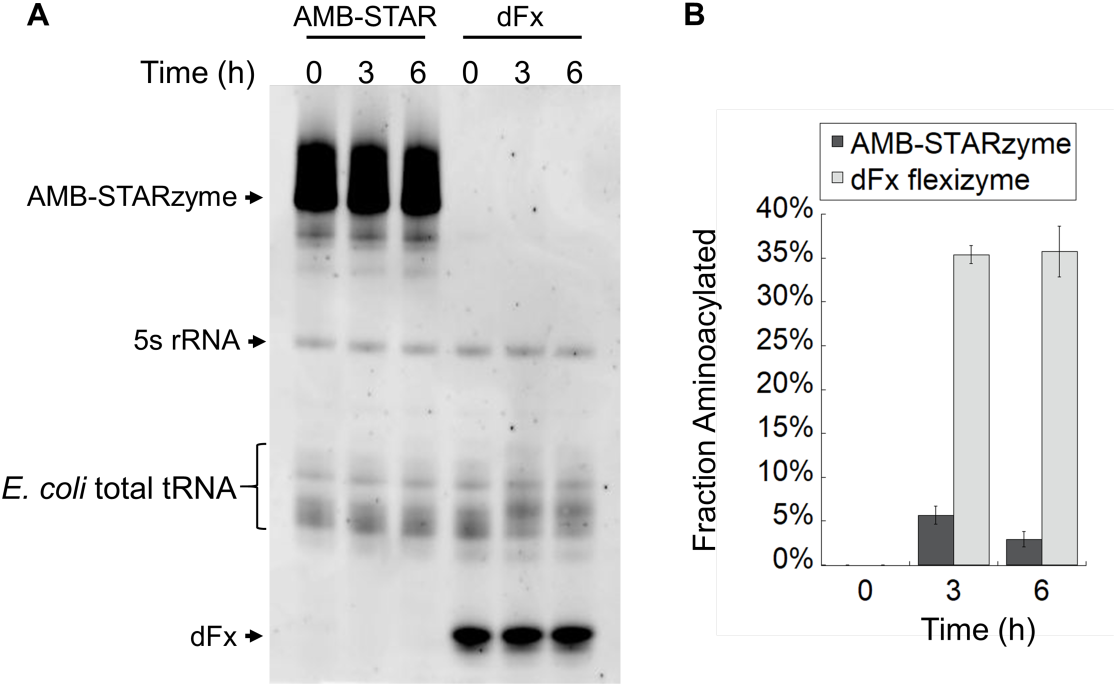
Testing the catalytic activity of AMB-STARzyme against *E. coli* total tRNA. **A.** A representative gel showing *E. coli* total tRNA aminoacylated by AMB-STARzyme or dFx over time. **B.** The calculated fraction of *E. coli* total tRNA aminoacylated by AMB-STARzyme (black) or dFx (grey).

### Using STARzyme to incorporate ncAA into proteins

To determine if the AMB-STARzyme can function in an *in vitro* translation system isolated from *E. coli*, the AMB-STARzyme/*M. jannaschii* tRNA^AMB^_CUA_ pair was introduced into the PURE *E. coli* translation system along with DBE- propargyl glycine in a single pot reaction. Translation reactions were carried out in the absence of release factor 1 (RF1) to suppress termination at the UAG Amber STOP codon. Control experiments indicated that the activated amino acid substrate is stable up to 24 h under these conditions. A mutant DHFR DNA template was constructed in which the proline 105 codon was replaced with the amber stop codon UAG. This site was chosen because it could be readily analyzed by MALDI-TOF mass spectrometry. AMB-STARzyme, *M. jannaschii* tRNA^AMB^_CUA_ and DBE-alkynyl amino acid were added into the system, along with the mutant DHFR DNA template.

In a control experiment, we found that adding only the P105TAG-DHFR DNA template in the absence of STARzyme, *M. jannaschii* tRNA^AMB^_CUA_ and DBE-propargyl glycine amino acid can produce low levels full-length DHFR protein, indicating that endogenous *E. coli* tRNAs are suppressing the UAG stop codon. This observation is consistent with a previous report noting that tyrosine, phenylalanine, tryptophan, and glutamine can be misincorporated at amber codons in absence of the release factor 1. Therefore, PAGE analysis alone could not unambiguously establish whether the amino acid incorporated into the polypeptide chain is the propargyl glycine charged onto tRNA^AMB^_CUA_ by the AMB-STARzyme (**Figure S5**).

To resolve this question, incorporation of the ncAA at the amber STOP codon site was evaluated by separating the protein products by SDS-PAGE, excising the target bands, digesting them with trypsin, and analyzing the peptide fragments by MALDI-TOF mass spectrometry. The WT sequence of the tryptic fragment containing position 105 is VYEQFL**P**K, with a calculated m/z of 1023.551 (singly charged). Indeed, using WT-DHFR DNA template yielded a peak with m/z of 1023.089, corresponding to the WT tryptic fragment (**Figure 7**). The recoded ncAA-containing fragment has the sequence of VYEQFL**X**K, with a calculated m/z of 1021.535 (singly charged). When the DHFR DNA template containing a TAG codon at a position corresponding to amino acid 105 was used along with the AMB-STARzyme, *M. jannaschii* tRNA^AMB^_CUA_ and DBE-propargyl glycine, a peak at /z of 1021.494 was clearly observed, indicating that the proline from the WT protein was replaced with a propargyl glycine residue at position 105. To further characterize the protein products, we used LC-MS/MS because it allows more sample injection, peptide separation, and low abundance ion detection. In the suppression experiment, the doubly charged precursor ion at 511.27 Da, corresponding to VYEQFL**X-2**K protein, was selected and fragmented **(Figure S6)**. The mass of the fragment ions could be unambiguously assigned, confirming the site-specific incorporation of propargyl alanine at position 105 **(Figure 7)**.

**Figure 7.**
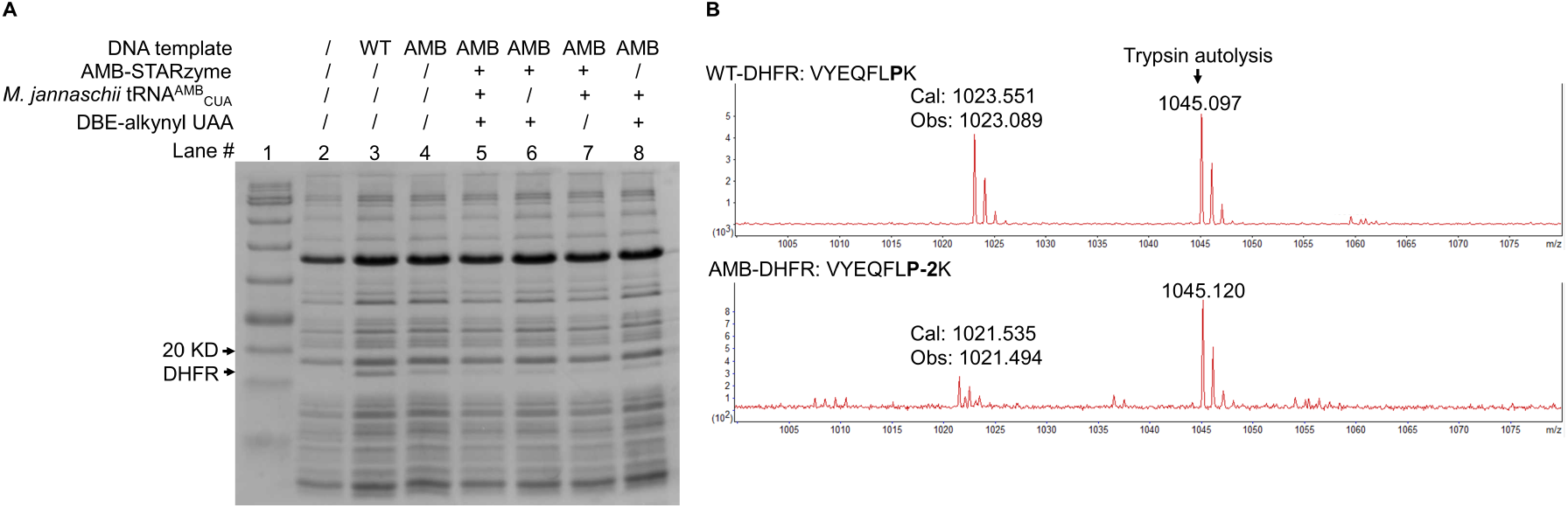
Coupling of AMB-STARzyme with in vitro translation. **A.** SDS-PAGE gel showing the in vitro translation with various conditions. Lane 1 is protein ladder and lane 2 is negative control with no DNA template added. **B.** MALDI-TOF detection of the translated products from lane 3 and lane 5. Calc, calculated; Obs, observed.

## Discussion

Here we describe the rational engineering of a ribozyme with aa-tRNA charging activity by fusing a tRNA binding module to a catalytic module. The molecular design is based on a T-box, a bacterial tRNA-binding aptamer, and laboratory-evolved biorthogonal aminoacyl transferase ribozyme. T-boxes recognize specific tRNAs through several interactions, which can be separated into a domain that base-pairs with the tRNA anticodon loop and that recognizes the elbow region of the tRNA through a quarternary interaction, and a domain that senses whether the tRNA is aminoacylated. We used the first domain as an aptamer-like module that recognizes a specific tRNA and fused it to the previously developed flexizyme which indiscriminately aminoacylate tRNAs. The fused ribozyme retained the catalytic activity of the flexizyme but gained tRNA specificity, allowing cognate tRNA charging is presence of other tRNAs.

The STARzyme was able to charge its cognate tRNA in a one-pot *E. coli* translation system, thereby directing a ncAA to a specific site in a protein. This reaction occurred in the presence of all the *E. coli* tRNAs and other translation components, suggesting it has sufficient tRNA specificity to function in a commercial translation system and, potentially, *in vivo*. As the flexizyme amino acid substrate is activated using a dinitrobenzyl ester leaving group, it cannot be recognized by cellular tRNA synthetases, and this property limits incorporation of the ncAA to the site specific location within the target protein.

### Towards the development of a universal tRNA synthetase

One benefit of this approach is that it is possible to rapidly devise variants that can in principle accommodate any tRNA and any amino acid by replacing the T-box module with a module specific for an alternate tRNA or supplying a different chemically activated amino acid. Here, we demonstrated that the *G. kaustophilus glyQS* T-box riboswitch in the initial design could be readily substituted with the *B. subtilis tyrS* T-box riboswitch (Grundy and Henkin 1993). The specificity for the anticodon of the tRNA can be altered using a much smaller change by mutating the specifier nucleotides in the T-box region (pink in **Fig 1).** Because the flexizyme has low specificity for the amino acid side chain (Coronado, Ngo et al. 2022), and thus the tRNA substrate can be charged with a large array of amino acids, including isotopically-labeled variants (e.g., ^15^N-Gly replacing ^14^N-Gly) and close analogues of natural amino acids, such as fluorinated amino acids. Near-identical analogues of natural amino acids that may be substrates of endogenous synthetase enzymes and would thus not allow site-specific incorporation of these variants in polypeptides *in vivo*. Our results suggest that the STARzyme may be a universal tRNA synthetase that will allow, via a biorthogonal chemistry, to charge tRNAs such that even chemically equivalent, but isotopically distinct, amino acids can be incorporated into polypeptides at specific locations using standard translation machinery. We anticipate that STARzymes will find applications in many areas of synthetic, chemical, and structural biology.

### Rational design of modular ribozymes from riboswitches

In the over 40 years since catalytic RNAs were discovered, a wealth of functional RNAs both naturally occurring and artificially evolved, have been discovered. In addition, many of these have been structurally characterized. These provide a tremendous library of tools that can be used for creating multifuctional modular RNAs for use as tools in basic research and biotechnologies.

Riboswitches account for many of these potential modules, as they tightly bind key metabolites within cells. It is certainly possible that modern riboswitches are the descendants of ribozymes that carried out metabolism in an early RNA World. T-box riboswitches look like they have the potential to have once been catalytic because they not only bind the body of the tRNA but also interact with the CCA tail to assess the charged-state of the tRNA. As proteins took over these key roles, some of these riboswitch predecessors may lost catalytic function, but were retained and adapted for gene regulation in modern biology. Here, we demonstrate one mechanism to ‘reawaken’ an ancient role for RNA by providing a catalytic domain to a catalytically inert binding module.

## Acknowledgements

Pew Charitable Trust Innovation Fund, NASA ICAR, Dr. Pallavi Joshi, Leon, Rob

## ^1^Abbreviations

DBE: dinitrobenzyl ester
dFx: a flexizyme specific for 5-dinitrobenzyl ester-activated amino acids
ncAA: non-canonical amino acid
STARzyme: specific tRNA aminoacylating ribozyme
T-box: tRNA-specific riboswitch
aaRS: aminoacyl tRNA synthetase.

